# Spatial transcriptomic profiling of decalcified murine musculoskeletal samples via Xenium Prime 5K

**DOI:** 10.64898/2025.12.22.693132

**Authors:** Lindsey Lammlin, Huong X. Tran, Aanya Mohan, Michael D. Newton, Scarlet C. Howser, Stephen J. Redding, Christina A. Capobianco, Kurt D. Hankenson, Alexander J. Knights, Tristan Maerz

## Abstract

Successful generation of high-quality spatial transcriptomics data from murine musculoskeletal tissues has been impeded by the challenge of preserving RNA integrity through the harsh tissue processing steps required for histological sectioning. In particular, the need to thoroughly fix and decalcify mineralized tissues has proven problematic. We detail a comprehensive sample processing pipeline for three common murine musculoskeletal tissue samples, enabling high-quality transcript detection via imaging-based spatial transcriptomics using the Xenium Prime 5K platform from 10x Genomics. Our protocol outlines methodological details for transcardiac perfusion, fixation, decalcification, paraffin processing, and a sample co-embedding strategy facilitating anatomically consistent and simultaneous sectioning of multiple samples onto the spatial transcriptomics slide. Rigorous quality control demonstrates high-quality tissue-specific outcomes across intact knee joints, tibiae, and lumbar spines from adult mice. Our pipeline enabled 70-91% high-quality transcripts across synovium, meniscus, patellar tendon, articular cartilage, subchondral bone, cortical bone, bone marrow, muscle, fracture callus, and dorsal root ganglion tissues. The average number of detected transcripts varied markedly between tissue types – soft tissues such as synovium, patellar tendon, muscle, bone marrow, and callus exhibited ~200 – 400 transcript per cell; mineralized tissues such as subchondral bone, meniscus, and cortical bone exhibited ~ 13 – 150 transcripts per cell; highly active neuronal tissues such as dorsal root ganglion neurons yielded 750 – 1100 transcripts per cell. Canonical cell markers within each tissue confirmed successful identification and representation of key cell types.

Through rigorous sample quality assessment at multiple stages of processing, this protocol yields high-quality RNA transcript detection while preserving critical anatomical context and will serve as a valuable tool enabling spatial transcriptomic profiling of intact musculoskeletal tissue samples.

**Lay summary:** Spatial transcriptomics is a powerful scientific tool that characterizes genetic coding material (“transcripts”) of specific cell types in their native anatomical context. However, successfully applying this tool to musculoskeletal tissues has been challenging because preserving transcript integrity in these tissues requires additional care. The authors of this publication developed a specialized method for preparing tissue samples from mice that works with the spatial transcriptomics platform, 10x Genomics Xenium Prime 5K. This protocol improves sample quality while preserving high-quality genetic information in various mouse musculoskeletal tissues, making it easier to explore their functions.

**Graphical abstract:** 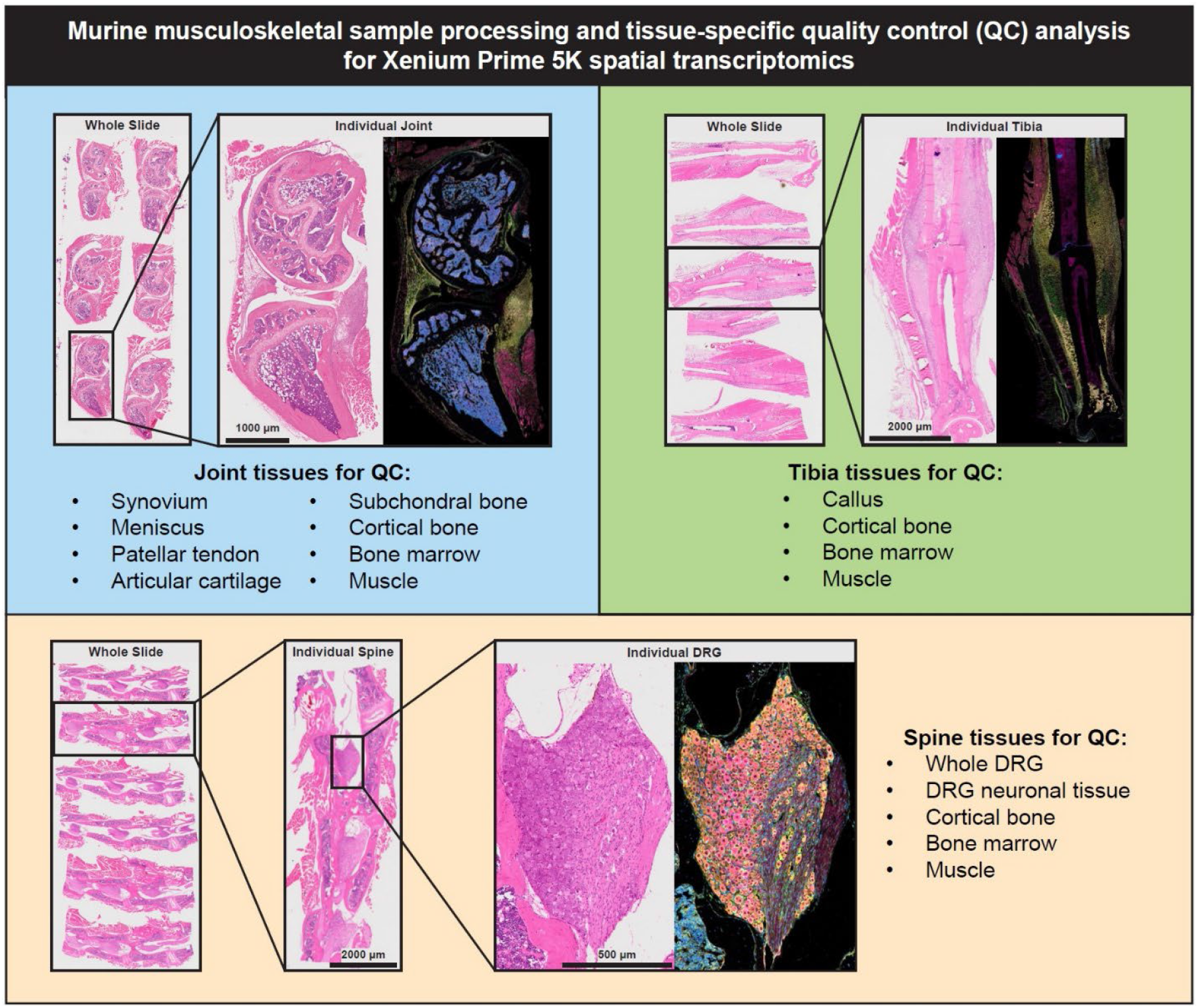

## Introduction

Spatial transcriptomics provides powerful insights into tissue biology, combining cell-specific gene expression with spatial information to interrogate complex tissue microenvironments in their intact state. In the context of musculoskeletal research, the application of spatial transcriptomics has been especially challenging due to the heterogeneous, extracellular matrix-rich composition of musculoskeletal tissues and sites^1^. Mineralized tissues pose a significant challenge during histological processing for spatial transcriptomics, as decalcification compromises RNA integrity. However, adequate decalcification is critical for preservation of tissue integrity during histological sectioning and the lengthy processing associated with spatial transcriptomics. At the cellular level, preserving the quality and native phenotype of cell types prone to rapid de-differentiation, apoptosis, or necrosis after euthanasia (for instance neurons^2,3^) represents an additional challenge. Here, we present a protocol for preparing intact, formalin-fixed, paraffin embedded (FFPE) murine musculoskeletal samples for spatial transcriptomic profiling using the Xenium Prime 5K platform from 10x Genomics (**Fig. 1**). Importantly, our protocol preserves the integrity of both mineralized and non-mineralized musculoskeletal tissues throughout the body as well as the dorsal root ganglion (DRG), a key orchestrator of musculoskeletal-related pain signaling. Our quantitative evaluation of sample quality and the resulting spatial transcriptomic data demonstrate that this pipeline enables the capture of high-quality transcripts at the single-cell level to enable identification and phenotyping of a wide range of cellular subsets in murine knee joints, tibiae, and spines.

**Figure 1.**
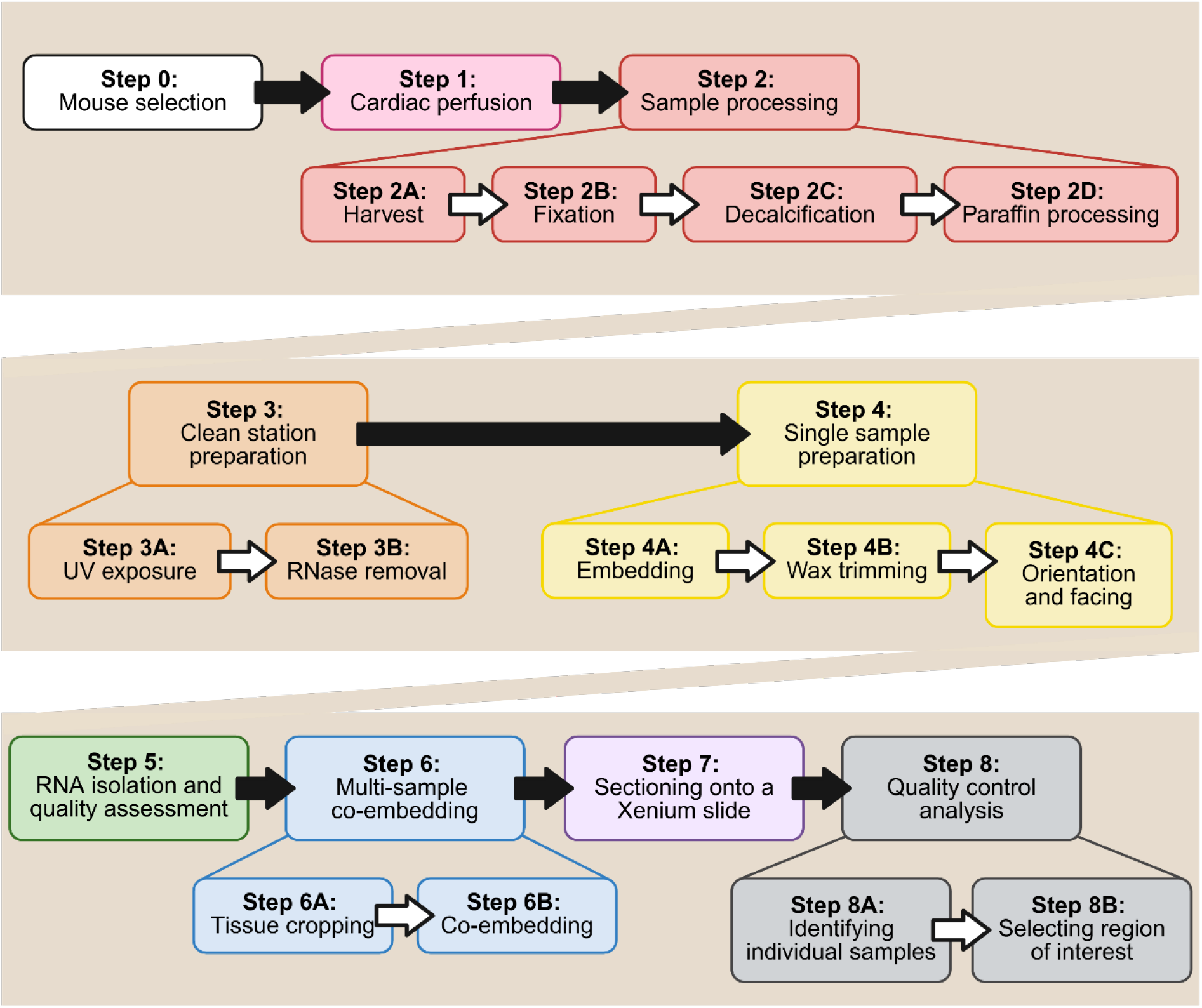
Protocol overview for 10x Genomics Xenium Prime 5K spatial transcriptomics of multiple co-embedded formalin-fixed, paraffin-embedded samples.

## Materials and Methods

### Step 0: Mice

All samples are from studies approved by the University of Michigan Institutional Animal Care and Use Committee. The workflow is optimized for musculoskeletal tissues from mice aged 12-18 weeks. We employed knee joints and lumbar spines (L3-L5) from naïve mice or mice that underwent non-invasive anterior cruciate ligament rupture (ACLR), and tibiae in the context of long-bone fracture (**Table 1**). However, this protocol can be applied to other murine musculoskeletal sites. A total of *n*=22 mice were utilized, with paired knee joints and spines taken from *n*=4 naïve mice and *n*=12 ACLR mice, and fractured tibiae from *n*=6 mice.

**Table 1.** List of samples included in quality assessments.

| Slide | # of Samples | Sample Conditions | Total # of Samples |
| --- | --- | --- | --- |
| Joint #1 | 6 | Healthy (x2), ACLR (x4) | Joints = 16 |
| Joint #2 | 6 | Healthy (x2), ACLR (x4) |  |
| Joint #3 | 4 | ACLR (x4) |  |
| Tibia #1 | 6 | Fracture (x6) | Tibiae = 6 |
| Spine #1 | 4 | ACLR (x4) | Spines = 16 |
| Spine #2 | 4 | Healthy (x2), ACLR (x2) |  |
| Spine #3 | 4 | Healthy (x2), ACLR (x2) |  |
| Spine #4 | 4 | ACLR (x4) |  |

### Step 1: Transcardiac perfusion

Systemic perfusion with fixative is critical to ensure rapid and widespread stabilization of RNA integrity. Deep anesthesia was induced using isoflurane (5% induction, 3% maintenance). Mice were secured in the supine position, the thoracic cavity was opened, and a 26G butterfly needle was inserted into the left ventricle. The mouse was perfused with 20 mL of cold PBS, followed by 50 mL of cold Z-fix Zinc Formalin Fixative (Cancer Diagnostics; SKU: 175) at a flow rate of 200 mL/hr. The total perfusion process took ~40 minutes per mouse from the time of anesthesia induction.

### Step 2: Sample processing: harvest, fixation, decalcification, and paraffin processing

Following perfusion, samples were harvested. For knee joints, the hindlimb was skinned, disarticulated at the femoroacetabular joint, and the hindpaw was removed. For fractured tibiae, the distal limb was skinned, disarticulated at the tibiofemoral joint, and the hindpaw was removed. For spines, the dorsal surface was skinned, and the lumbar spinal column was removed. Samples were fixed for an additional 48 hours in 10 mL Z-fix while nutating at 4ºC. Samples were decalcified within one week of harvest with 80% StatLab™ Immunocal (Fisher Scientific; STL141432), diluted in PBS, while nutating at 4ºC, optimizing timing based on bone density (joints and spines: 16 hours; tibiae: 18 hours). After decalcification, samples were stored in 70% ethanol at 4ºC until paraffin processing. Samples were paraffin-processed using Program 1 (joints and spines) or Program 2 (tibiae) (**Table 2**).

**Table 2.** Paraffin processing details for Programs 1 and 2 which are available at the Michigan Integrative Musculoskeletal Health Center’s Structure, Composition, and Histology Core. Steps were carried out under a vacuum at ambient temperature (Steps 1-10) or 60ºC (Steps 11-13).

| Step of Processing | Solution | Time (Program 1) | Time (Program 2) |
| --- | --- | --- | --- |
| 1 | 70% ethanol | 1 hr | 2 hr |
| 2 | 80% ethanol | 1 hr | 2 hr |
| 3-4 | 95% ethanol | 2x (1 hr) | 2x (2 hr) |
| 5-7 | 100% ethanol | 3x (1 hr) | 3x (2 hr) |
| 8-10 | Xylene | 3x (1 hr) | 3x (2 hr) |
| 11-13 | Paraffin | 3x (1 hr) | 3x (2 hr) |

### Step 3: Clean station preparation for paraffin sectioning

To maximize RNA integrity, all paraffin sectioning was performed in a clean, dead-air box (Mystaire; MY-DB48). Multiple precautions were taken to minimize the presence of RNA-compromising contaminants. Prior to usage, the workstation was exposed to UV light for 15 minutes. All sectioning tools and working surfaces inside the working station, including the microtome, and slide warmer, were cleaned with RNaseZap (ThermoFisher; AM9780). Autoclaved water was used in the sample collection water bath, which was set to 50ºC. The slide warmer was set to 37ºC. The operator wore sterile gloves and a mask.

### Step 4: Single sample embedding, orienting, and trimming

Samples were individually embedded in large tissue embedding molds (Epredia; 64-010-18). Excess muscle was trimmed to allow the sample to lie flat on the desired sectioning surface. Spine samples were further trimmed to remove non-applicable spinal levels via transverse cuts through the L3/L4 intervertebral disc, and midway through the L6 vertebral body, preserving the knee-innervating L3-L5 DRGs. Spines were further bisected in the sagittal plane to generate a left and right sample, enabling separate localization of left/right knee-innervating DRGs. For this work, only ipsilateral (ACLR side) spine samples were utilized. Samples were then placed in a melted paraffin wax bath at 60ºC for 10 minutes to allow complete paraffin infiltration before embedding. For knee joints, the medial side was oriented downwards (i.e., for sectioning into the medial compartment); tibiae were oriented posterior side down, and spines with the medial, bisected side down. Paraffin-embedded samples were stored at 4ºC until individual trimming and orienting.

During individual sample trimming, tissue sections were collected for RNA quality assessment, and samples were faced with a consistent anatomic orientation and sectioning plane. Excess paraffin was cut from each paraffin block to minimize paraffin content during RNA isolation and quality assessment, as we have observed that excess paraffin is associated with lower RNA integrity number (RIN) (**Fig. 2**). Five 20-µm sections were collected into an RNase-free tube and stored at 4ºC (**Fig. 2**). Trimming was halted ~150 µm before the desired depth, i.e. anatomic location, to allow for final fine-tuning of orientation and depth with co-embedded samples. A paraffin cap was placed over the exposed sample to prevent desiccation or damage, and samples were stored at 4ºC.

**Figure 2.**
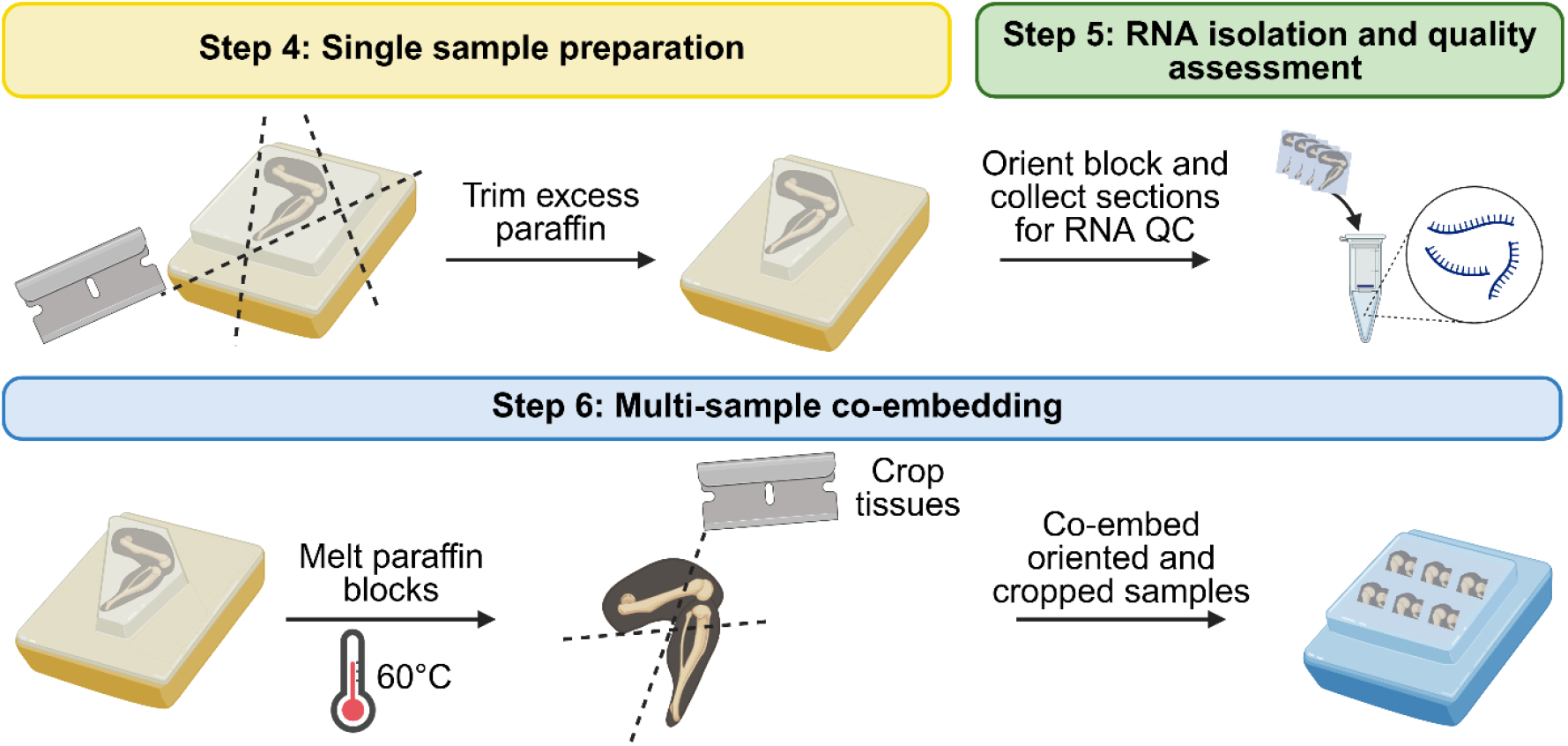
Histological preparation workflow.

### Step 5: RNA isolation and quality assessment

RNA was isolated from tissue sections using the RNeasy FFPE isolation kit (Qiagen; 73504). Due to the large amount of tissue and paraffin material, we doubled reagent volumes recommended in the manufacturer’s protocol to effectively facilitate RNA purification. RNA quality was evaluated using the Agilent Bioanalyzer to yield RNA integrity number (RIN, scale from 1-10) and fragmentation of RNA (DV200, %) (**Table 3**). For this study, RNA integrity from joint tissues was deemed representative of paired spines. For the Xenium platform, although there is no minimum RIN recommendation, 10x Genomics recommends a DV200 above 30% to ensure high-quality data^4^.

**Table 3.** RNA quality assessment of FFPE sections.

|  | Joints | Tibiae |
| --- | --- | --- |
| **RIN** | 4.4 ± 0.7 | 3.3 ± 0.4 |
| **DV200** | 74.68% ± 6.08% | 77.85% ± 3.27% |

### Step 6: Multi-sample co-embedding

Given the high cost of spatial transcriptomics, we aimed to maximize the number of samples that could be processed on each Xenium slide using a co-embedding strategy. Samples were trimmed macroscopically with a razor blade to remove excess tissues (**Fig. 2**). All cropped tissues were positioned in the small co-embedding mold (10 x 24 x 5 mm, TedPella; 27276-2) in a predetermined layout. The trimmed side of each sample was placed downwards into the mold on a thin coat of melted paraffin wax. The mold was briefly placed on a cold plate to secure the samples, followed by complete filling with paraffin wax. The co-embedded block was stored at 4ºC until final sectioning. Co-embedding up to six knees or six tibiae was achieved with a small tissue mold, yielding a block from which a single section could be cut to fit into the capture area of a Xenium slide (10.45 x 22.45 mm) (**Fig. 2**). Due to the small DRG region of interest (ROI), spines were individually embedded, and tissue sections from separate blocks were consecutively placed onto the Xenium capture area to ensure proper orientation of each sample.

### Step 7: Sectioning onto a Xenium slide

The Xenium slide was removed from storage and equilibrated to room temperature for at least 30 minutes. Final fine-trimming of the co-embedded samples was performed at 5-µm increments. As with standard paraffin histology, tissue blocks were iced and hydrated prior to collecting the final Xenium section. Once all samples were in-view/in-plane at the desired ROI, a final tissue section was collected (5-µm thickness) onto the Xenium slide, taking care to overlay the section onto the capture area without covering the fiducials, consistent with 10x Genomics protocol^5^. A wax cap was placed over the exposed tissue surface, and the block was stored at 4ºC as described in *Step 4*. The Xenium slide was fully dried on a slide warmer for ~35 minutes at 37ºC. According to the 10x Genomics protocol, the Xenium slide with tissue sections can be stored in a slide box containing a desiccant at room temperature for up to 14 days before processing^5^; ideally though, it should be submitted for processing as soon as possible. We utilized the University of Michigan Advanced Genomics Core (UM-AGC) for all post-section collection processing of the Xenium slide utilizing the standard pipeline and reagents from 10x Genomics for the Xenium Prime 5K platform.

### Step 8: Data quality control (QC)

Xenium Analyzer (v3.3.0.1) was used for imaging and transcript detection. Xenium Ranger (v3.3.0.1) was used for raw data processing. Large cell segmentation was utilized for spine samples^6^ (**Supplement**). Xenium Explorer (v3.2.0) was used for sample and tissue selection, transcript overlay, and image export. R (v4.5.0) and RStudio (v2024.12.1) were used for object processing and QC metric calculations. The following R packages were used for QC: Seurat (v5.3.0)^7^, tidyverse (v2.0.0)^8^, arrow (v21.0.0)^9^, and Matrix (v1.7-3)^10^.

### 1. Xenium gene probe panel

We employed the Xenium Prime 5K Mouse Pan Tissue & Pathways Panel, targeting 5006 genes with an additional 100-gene custom panel, focused on genes relevant to musculoskeletal research. Additionally, we utilized the multimodal cell segmentation, which employs a panel of antibodies targeting the cell membrane (E-cadherin, ATPase1a1, CD45), cytoplasmic RNA (18s), cytoplasmic protein (αSMA, Vimentin), and nuclei (DAPI), to increase the fidelity of cell segmentation^11^. Although this step is optional, we highly recommend it for post-processing cell identification and quality control analysis.

### 2 .ROI selection

#### a) Identifying individual samples

Individual samples on the Xenium slide may be identified during Xenium Analyzer processing, or in Xenium Explorer after processing the entire slide as one sample. Additional information on individual sample identification during processing can be found in the **Supplement**. Incorrect or incomplete sample identification using the Xenium Analyzer prior to probe hybridization is problematic as only the selected regions will be processed for transcript detection rather than the entire slide, which could result in missing data. Therefore, our recommendation is to process the entire slide and identify samples in Xenium Explorer, followed by individual sample data separation in R.

#### b) Performing tissue outlines by sample

Following the Xenium assay, Hematoxylin and Eosin (H&E) staining was performed by the UM-AGC to assess tissue morphology. Transcript and cell segmentation information is embedded in the Xenium morphology image; therefore, aligning the H&E and Xenium morphology images allows for accurate identification of ROIs (from the H&E) and extraction of transcript information within those ROIs (from Xenium morphology image) for downstream analysis. The H&E image for the corresponding slide was imported into Xenium Explorer and aligned by placing ~200 registration points per sample. DAPI-derived nuclei boundaries were used to verify alignment with nuclei on the H&E image. If the alignment was unsatisfactory, additional points were added. Following alignment, both the H&E and Xenium morphology images were used for ROI selection. Outlines were drawn using the selection tool in Xenium Explorer. Coordinates were extracted and used to computationally subset cells from these regions. Tissue regions defined in each sample are shown with selected gene transcript overlays in **Figures 4** (joints), **5** (tibiae), and **6** (spines).

#### c) Segmentation of neuronal tissue within spinal sections

Whole DRG of spinal tissues were outlined and further segmented to identify neuronal regions of the DRG for analysis. This extra step of QC was necessary to confirm successful extension of this pipeline to the highly neuron-rich tissue, as neurons are among the first cell types to die upon euthanasia^2,3^. Therefore, separate evaluation of the quality of neuronal DRG tissue was performed to ensure quality within these sensitive regions.

### 3) Quality control metrics

Quality control metrics were generated in R to evaluate probe hybridization, transcript detection, and cell segmentation across tissues.

#### a) Probe Hybridization

The Xenium Prime 5K assay includes built-in control probes to evaluate the specificity of the probe-binding assay and gene decoding algorithm, which can be used to calculate the False Discovery Rate (FDR)^12^, identifying off-target binding events. The values are expected to be close to 0% in a robust panel design with high gene probe specificity^12^.

#### b) Transcript Quality

The Xenium Analyzer pipeline calculates a Q-value for each transcript, estimating the probability of a transcript being incorrectly assigned to a gene during probe hybridization. Transcripts with a Q-value ≥ 20 are considered high-quality^13^. Transcript quality scores were extracted from the *transcripts*.*parquet* file. Transcripts from cells within each tissue region were extracted. The percentage of total transcripts with a Q-value ≥ 20 and the average number of high-quality transcripts per cell were calculated for each tissue, by sample.

#### c) Nucleus Count

To examine the cell segmentation quality, we evaluated the nucleus count per cell, expecting most cells in a tissue to be mononucleated. The number of nuclei per cell was extracted from the *cells*.*parquet* file and then written into the metadata using the corresponding cell ID. The percentage of mononuclear cells was calculated for each tissue, by sample.

### Step 9: Tips and tricks

We highly recommend that users read the additional troubleshooting tips in the **Supplement**.

## Results

Our protocol successfully yielded high-quality spatial transcriptomics data in murine tissue from multiple musculoskeletal sites. Despite multiple rounds of tissue staining and probe hybridization, imaging of immunofluorescent and H&E-stained samples confirmed intact tissue sections, with only rare instances of histological artifacts such as tissue tearing or folding. Assessment of RNA quality was performed using RIN, which measures the presence and ratio of ribosomal RNA, and DV200, which measures the percentage of intact RNA fragments >200 nucleotides in length. DV200 is superior for assessing transcripts in fixed samples and better reflects usable fragments for sequencing. Our knee joint and tibia FFPE sections had RIN values ranging from 3-5 and DV200 values ranging from 70-80% (**Table 3**). QC outcomes related to built-in negative controls showed near-zero off-target probe binding, with FDR ranging from 0.1-0.7% (**Table 4**), confirming a high degree of assay specificity.

**Table 4.** Mean False Discovery Rate (FDR, %) of negative control probe binding per slide.

| Slide | Mean FDR (%) ± Margin of Error |
| --- | --- |
| Joint #1 | 0.325 ± 0.038 |
| Joint #2 | 0.302 ± 0.039 |
| Joint #3 | 0.556 ± 0.15 |
| Tibia #1 | 0.7 ± 0.131 |
| Spine #1 | 0.187 ± 0.027 |
| Spine #2 | 0.255 ± 0.057 |
| Spine #3 | 0.315 ± 0.072 |
| Spine #4 | 0.3 ± 0.045 |

We observed a high mean proportion of high-quality transcripts across all analyzed tissues, ranging from 73.6-88.7% (SEMs range 0.303-0.736%), indicating that our sample processing yielded RNA sufficient for the detection of intact, high-quality transcripts (**Fig. 3A**). The neuronal portion of DRGs exhibited the lowest percentage of high-quality transcripts (mean = 73.6%), consistent with prior studies demonstrating more rapid RNA degradation in neurons^2^. The average number of transcripts per cell by tissue ranged from 13 to 1,118, with articular cartilage exhibiting the fewest transcripts per cell (sample mean = 34; SEM = 3.36), and DRG neuronal tissue with the highest (sample mean = 934, SEM = 40.71) (**Fig. 3B**). The mean proportion of cells with one nucleus ranged from 97.1-100.0% across all samples with the lower values in muscle (mean = 98.2%) and bone marrow (mean = 98.6%), in which multinucleated cells are more common, and whole DRG (mean = 97.1%) and neuronal DRG tissue (mean = 97.4%), likely reflecting a small percentage of cases where neuronal cell boundaries encompass the nuclei of surrounding support cells such as Schwann and glial cells (**Fig. 3C**).

**Figure 3.**
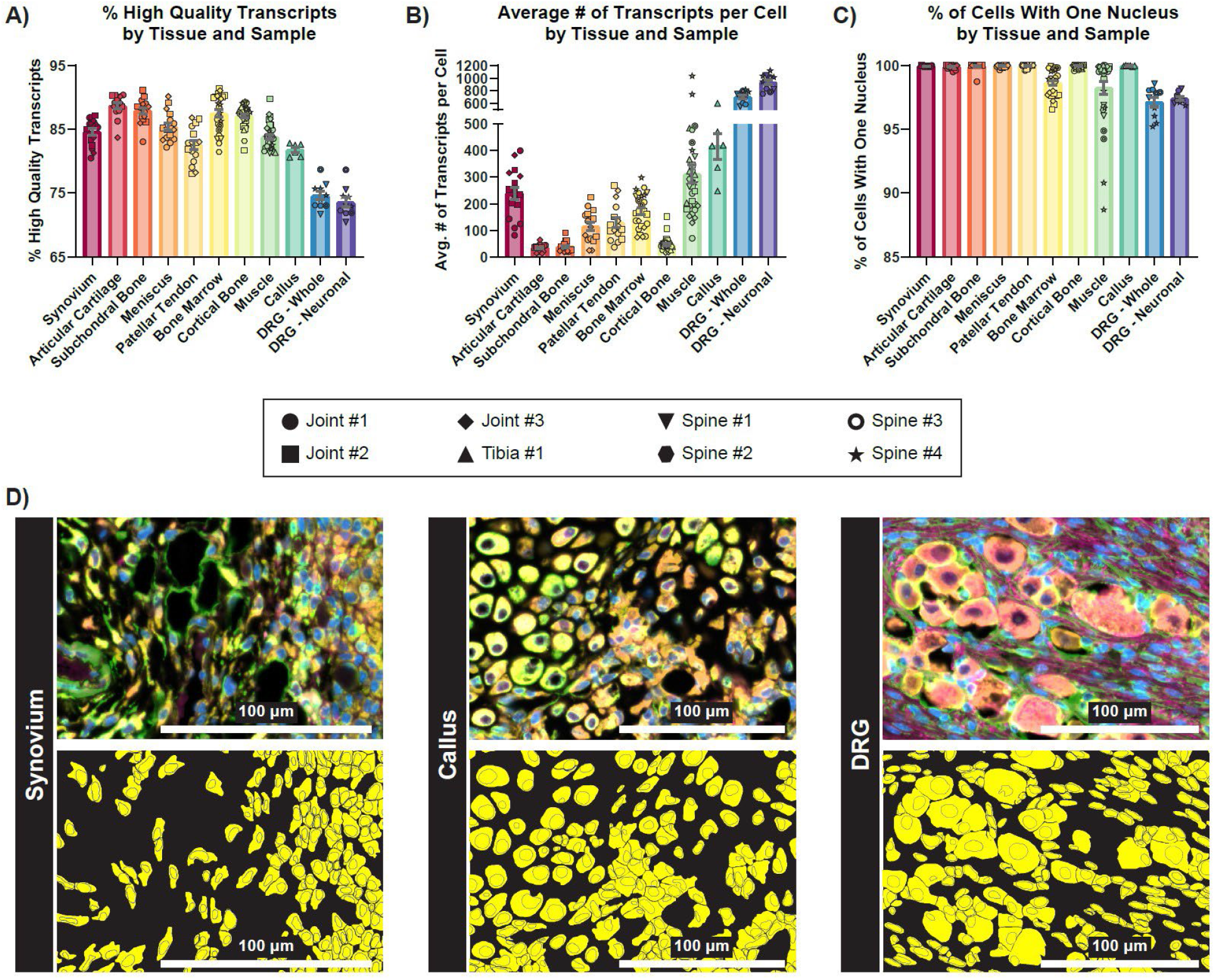
Quality control outcomes. A-C: Percentage of high-quality transcripts (Q-value ≥ 20) (A), average number of detected transcripts per cell (B), and percentage of cells with one nucleus (C), organized by tissue. Replicates represent individual sample sections, and data symbols denote the slide of origin. Error bars show mean ± SEM. (D) Representative examples of immunofluorescent imaging from the multimodal cell segmentation antibody cocktail (top) and from the downstream cell segmentation output showing individual cell outlines and their nuclei (bottom), illustrated in synovium, tibial callus, and DRG tissue. Scale bars: 100 µm.

Implementation of the optional 10x Genomics multimodal cell segmentation resulted in accurate cell boundary and nucleus segmentation, even in regions of high cell density and varying cell sizes (**Fig. 3D**). Our combined adaptation of the 10x large-cell segmentation model with standard segmentation resulted in robust identification of large neurons and small non-neuronal cells in the DRG (**Fig. 3D**). Representative transcript feature plots showed detection of canonical markers for key cell types at the single-cell level across a variety of tissues in knee joints (**Fig. 4**), tibiae (**Fig. 5**), and spines (**Fig. 6**). Even in tissues with low transcript counts, key cell markers such as *Runx2* in subchondral bone, *Sox9* in articular cartilage, and *Sost* in cortical bone were broadly detectable. In DRG neurons from spines, we detected key neuronal subset-specific markers (e.g. *Scn10a* enriched in nociceptors, *Nefh* enriched in neurofilament-positive, large-diameter neurons including mechanoreceptors and proprioceptors), in addition to markers of neuronal peripheral cells (e.g. *Fabp7* marking glial cells; *Mpz* marking Schwann cells) and nerve root-resident stromal and immune cells.

**Figure 4.**
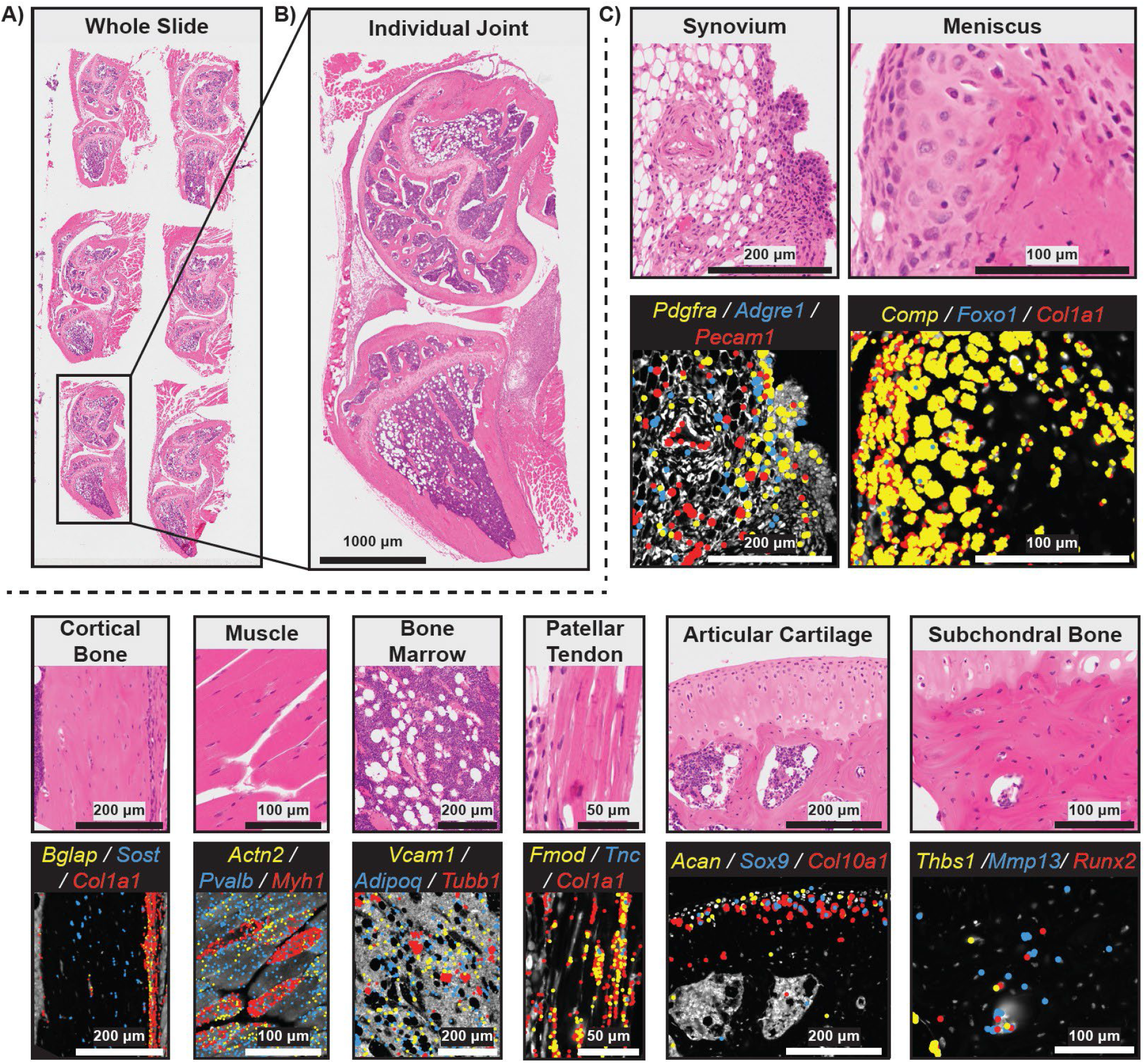
Representative example of a Xenium experiment with six healthy and injured murine knee joints. (A) H&E image of whole Xenium 5K slide. (B) H&E image of individual joint sample. (C) Tissue zooms with select gene overlays relevant to prominent cell types found in each joint tissue. Scale bars are shown on each image.

**Figure 5.**
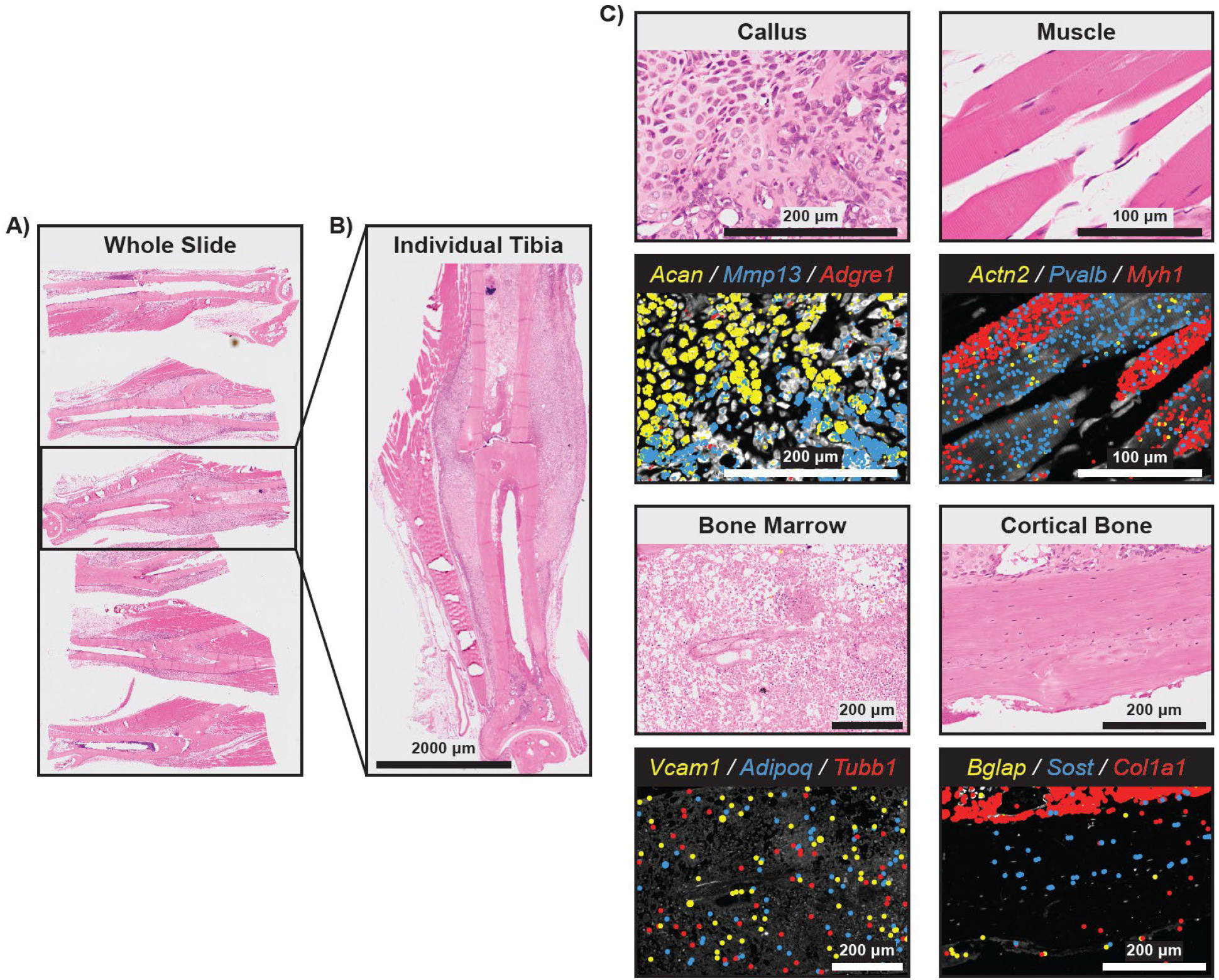
Representative example of a Xenium experiment with six fractured murine tibiae. (A) H&E image of whole Xenium 5K slide. (B) H&E image of individual tibia sample. (C) Tissue zooms with select gene overlays relevant to prominent cell types found in each tibial tissue. Scale bars are shown on each image.

**Figure 6.**
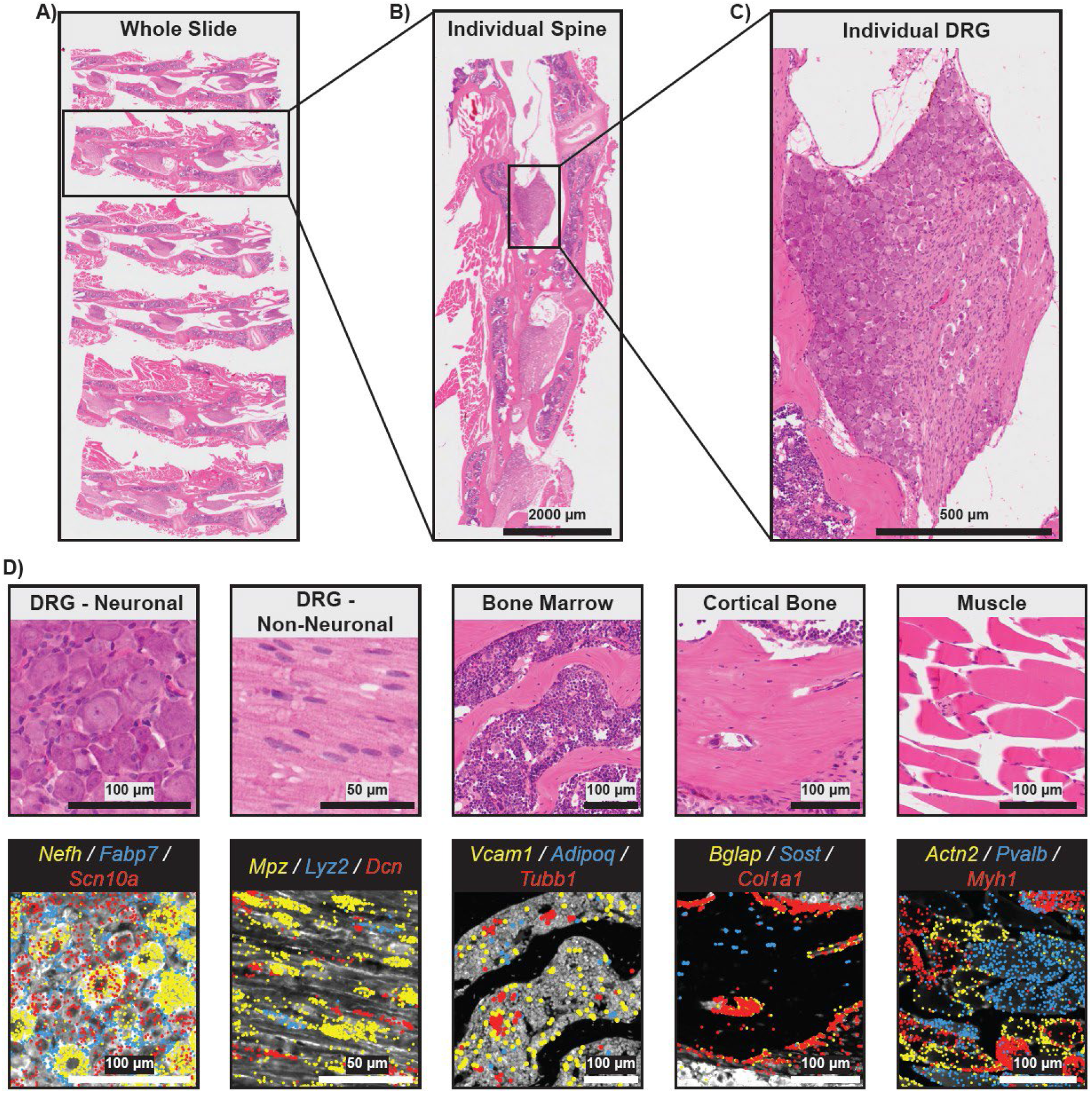
Representative example of a Xenium experiment with six murine lumbar spines (L3-L5). (A) H&E image of whole Xenium 5K slide. (B) H&E image of an individual spine sample. (C) H&E image of an individual DRG sample. (D) Tissue zooms with select gene overlays relevant to prominent cell types found in each spinal tissue. Scale bars are shown on each image.

## Discussion

The application of spatial transcriptomics to musculoskeletal research has been limited because the harsh processing required for FFPE musculoskeletal samples compromises RNA integrity. In this protocol we successfully surmount tissue processing challenges to generate high-quality spatial transcriptomics data from multiple murine musculoskeletal sites: knee joints, tibiae, and spines. During protocol development, we established that rapid sample fixation via transcardiac perfusion is a key step in preserving RNA quality. The subsequent fixation step further ensures complete RNA stabilization. We also observed that although the formic acid-based Immunocal is a stronger, more rapid decalcification agent than traditional ethylene diamine tetra-acetic acid, it enabled superior RNA quality (data not shown).

Imaging-based spatial transcriptomics technologies, including Xenium Prime 5K, generally yield fewer detected transcripts per cell compared to sequencing-based approaches, such as Visium^1^. Therefore, in tissues with low cellularity, such as articular cartilage, maximizing RNA quality and quantity is of utmost importance. However, it should be noted that the low transcript numbers in articular cartilage, bone, and meniscus are also directly related to the design of the assay, as gene probes related to chondrocyte biology are underrepresented in the Mouse Pan Tissue & Pathways Panel. Our additional custom 100-gene panel, specific to musculoskeletal applications, contributed to increased numbers of transcripts per cell. A limitation of our data is the absence of information regarding the intervertebral disc (IVD), another musculoskeletal tissue with low vascularity that may be prone to poor RNA quality. We also did not compare FFPE to cryohistology, as our goal was to generate samples amenable to spatial transcriptomics and our standard FFPE stains and analyses.

Through strict, multi-stage quality control, we have demonstrated that this protocol enables the generation of high-quality spatial transcriptomics data in murine musculoskeletal samples.

## Funding

LL was supported by funding from the National Science Foundation Graduate Research Fellowship grant (DGE 2241144).

CC was supported by funding from the National Institutes of Health (T32TR004371, F31AR085942).

KDH was supported by funding from the National Institutes of Health (R01AR066028).

AJK was supported by funding from the National Institutes of Health (R00AR081894) and the Arthritis National Research Foundation (1469667).

TM was supported by the National Institute of Arthritis and Musculoskeletal and Skin Diseases (NIAMS) of the National Institutes of Health (R01AR080035, UC2 AR082186).

KDH and TM were supported by funding from the Department of Defense Congressionally Directed Medical Research Programs (CDMRP) (HT94252310327).

Michigan Integrative Musculoskeletal Health Core Center’s (MiMHC) Structure, Composition, and Histology Core (NIAMS P30 AR069620).

University of Michigan, Department of Orthopaedic Surgery.

## Author contributions

Conceptualization: LL, HXT, AJK,TM

Methodology: LL, HXT, AM, MDN, SCH, SJR, AJK

Investigation: LL, HXT, AM, MDN, SCH, SJR,

CC Visualization: LL, HXT, AM

Funding acquisition: LL, CC, KDH, AJK, TM

Writing – original draft: LL, HXT, AM

Writing – review & editing: LL, HXT, AM, MDN, KDH, AJK, TM

## Data and materials availability

Not applicable.

