## Supplement for "Spatial transcriptomic profiling of decalcified murine musculoskeletal samples via Xenium Prime 5K"

### Tips and tricks

*Evaluating the success of the transcardiac perfusion*

Successful perfusion is critical to the Xenium workflow as rapid fixation is necessary for preservation of tissue and RNA quality[14,15]. In these studies, successful perfusion was indicated by one or all of the following observations: liver and lung clearing, muscle tremors/tail movement, and overall body stiffness. Clearing of the liver is often the first indicator of successful perfusion and is shortly followed by tremors, most obviously in the tail and legs. The lungs should remain flat and blanched. Regardless of the presence of these signs, proper perfusion should always result in body stiffness. At the conclusion of perfusion, the mouse should be extremely rigid, and there should not be any laxity or malleability at the limbs, neck, or spine, and the tail should extend stiffly without drooping due to gravity.

There are multiple errors arising during perfusion which can be addressed once recognized. The first problem that can occur is inflation of the lungs at the beginning of the perfusion. Immediate inflation of the lungs is likely due to the needle penetrating the right ventricle through the cardiac wall or pressure buildup without an outlet in the right atrium, and the perfusion may be saved by needle repositioning or right atrium snipping. Liver spotting (clearing) should occur within the first 2 minutes of the perfusion, and needle repositioning is required if this is not observed. At the conclusion of the fixative perfusion, if the mouse is malleable (ex. limbs and neck can be easily manipulated to articulate), then the needle was likely in the wrong position.

Noting the timing of successful perfusion indicators (liver clearing, tremors) and overall stiffness post-fixation allows the researcher to track sample quality throughout the Xenium workflow and identify sources of potential processing or technical issues, should the data not yield the expected quality comparing to the data presented in this study.

*Preserving sample integrity after paraffin processing and trimming*

If samples cannot be immediately embedded after paraffin processing, they can be stored at 4ºC until it is time for embedding. When samples are stored at 4ºC prior to embedding, they should be allowed to come to room temperature for 30 minutes prior to being put in the paraffin wax bath – this minimizes the amount of time required for the sample to sit in the paraffin wax bath. Placing a cold sample in the paraffin wax bath will require a longer amount of time to ensure complete wax infiltration, which may accelerate RNA degradation in the outer layers of tissue.

Once sample blocks have been trimmed (single samples or co-embedded samples), it is highly recommended to store them at 4ºC to mitigate RNA degradation after air exposure.

Placing a wax cap over the exposed tissue surface of the block can help prevent tissue desiccation and preserve tissue integrity during long term storage.

*Removing excess wax on individual samples prior to collecting tissue for RNA isolation*

While the RNeasy FFPE kit (Qiagen; 73504) for RNA isolation is designed specifically for this purpose, our iterative optimization studies (data not shown) showed that a large wax-to-tissue ratio will decrease the quality of RNA from the sample. To mitigate paraffin wax contamination in the sample, excess wax is rigorously trimmed from the individually embedded samples prior to collecting sections for RNA isolation.

*Troubleshooting poor tissue quality while sectioning I – decalcification*

Poor sectioning quality could be due to a number of issues, one of the most common being insufficient decalcification. X-ray imaging of samples can be used to confirm sufficient decalcification prior to paraffin processing. If the sample has been completely processed and embedded, and insufficient decalcification is suspected, the paraffin block can be x-rayed to confirm. Signs of insufficient decalcification include tissue tearing and a crunchy texture upon sectioning. For regular structural histology evaluations, samples would be de-paraffin processed, decalcified, and re-paraffin processed, or subjected to surface decalcification directly on the paraffin block. While these are potential solutions to insufficient decalcification, the researchers did not test and validate these methods for this Xenium workflow.

*Troubleshooting poor tissue quality while sectioning II – wax infiltration*

Another common cause of poor sectioning quality is insufficient wax infiltration into the sample, which is evidenced by a hole in an otherwise good quality tissue section. This can be amended by placing the exposed surface of the block in melted paraffin wax for a couple of minutes and then allowing it to completely solidify again. If the lack of paraffin wax infiltration is severe, the block can be completely melted down and re-embedded.

*Re-embedding co-embedded samples*

When sectioning the co-embedded sample block, it is possible that one or more of the samples are either slightly out of plane, differentially oriented, or not at the same depth as the others. In this case, the researcher may melt down and re-embed all samples in the block. However, if only one sample is differentially oriented or not sectioned into as far as the other samples, it is possible to remove just that sample from the block. To do this, allow the block to come to room temperature and heat up a razor blade, similar to that used for macroscopically cropping the samples. The problematic sample can then be cut from the co-embedded block, embedded individually, and the orientation and trimming depth can be fixed. Then, all the samples, including the re-adjusted sample and the co-embedded samples, can be melted down and re-embedded.

*Saving sections for later reference or staining*

When individually trimming samples, it is helpful to collect a few sections while approaching the desired ROI, especially the last section prior to finishing the trim. This will serve as a helpful reference to ensure that all samples are trimmed to the same depth. Additionally, these sections close to the desired ROI can also be used for other purposes, such as additional structural staining for histological assessment. Similarly, a few sections from the co-embedded block should also be collected prior to sectioning onto the Xenium slide. This will allow for additional immunofluorescent or structural staining that can be complementary to Xenium analysis.

*Identifying individual samples during Xenium Analyzer processing*

While we recommend identifying individual samples in Xenium Explorer, should the researcher choose to identify samples during Xenium Analyzer processing, we have provided some additional information to aid in this process. Before running the instrument, the individual sample boundaries are determined by the instrument operator. Each sample region is selected by a series of field of views (FOVs), each with an area of 3,520 x 2,960 pixels (0.2125 µm/px resolution), with 128 pixels of overlap to each adjacent FOV to ensure smooth stitching of the image and accurate transcript alignment[2]. The entire slide capturing area can be divided into around 19 x 33 FOVs that will be distributed to individual samples. Data generated will be specific for each selected sample region, which is convenient for downstream analysis. Since sample sections are close to one another, care must be taken to confine FOVs to a single sample, as tissue regions from one sample can be incorrectly grouped with a separate sample, or entire tissue segments may be cut off.

*Large cell segmentation of spine slides*

Neuronal cell bodies are not accurately segmented by the default cell segmentation algorithm used by Xenium Ranger, due to their larger size. To account for this, spine slides were re-segmented in Xenium Ranger (v4.0.1.1) with the large cell segmentation option *--segment-large-cells=true* activated. This preserves the default segmentation methods while adding an additional boundary-based segmentation layer calibrated for larger cells[6]. Note that at the time of writing, the large cell segmentation model provided by 10x is not trained on mouse neuron data – thus, while the large cell segmentation substantially improves segmentation, some manual correction may still be necessary to achieve rigorously accurate segmentations in all cells. Other options for cell segmentation have also been explored by other research groups, including Cellpose[16] and Baysor[17], among others, and these may also be considered as alternatives to the available 10x-provided methodologies.
